# Extracellular Pax6 regulates tangential Cajal-Retzius cell migration in the developing mouse neocortex

**DOI:** 10.1101/269480

**Authors:** H. Kaddour, E. Coppola, A. A. Di Nardo, A. Wizenmann, M. Volovitch, A. Prochiantz, A. Pierani

## Abstract

The embryonic mouse cortex displays a striking low caudo-medial and high rostro-lateral graded expression of the homeoprotein transcription factor Pax6, which presents both cell autonomous and direct non-cell autonomous activities. Through the genetic induction of anti-Pax6 single-chain antibody secretion, we have analyzed Pax6 non-cell autonomous activity on the migration of cortical hem- and septum-derived Cajal-Retzius (CR) neurons by live imaging of flat mount developing cerebral cortices. We observed that blocking extracellular Pax6 disrupts tangential CR cell migration patterns. We found a decrease in the distance travelled and changes both in directionality and in the depth at which CR cells migrate. Tracking of single CR cells in mutant cortices revealed that extracellular Pax6 neutralization enhances or reduces contact repulsion in medial and lateral regions, respectively. This study demonstrates that secreted Pax6 controls neuronal migration thus acting as a *bona fide* morphogen at an early stage of cerebral cortex development.

**Summary statement:** Cajal-Retzius cell distribution in the embryonic cortex participates in determining the size and positioning of cortical areas. Here, Kaddour et al. establish that the direct non-cell autonomous activity of the Pax6 transcription factor regulates Cajal-Retzius cell migration.

## Introduction

Glutamatergic Cajal-Retzius cells (CR cells) are a transient neuron population residing at the surface of the developing cerebral cortex (Marin-Padilla, 1998). They comprise molecularly distinct subtypes born in three different progenitor domains of the mouse cortical neuroepithelium: the cortical hem (CH), the septum (S) and the ventral pallium at pallial-subpallial boundary (Bielle et al., 2005; Yoshida et al., 2006; Tissir et al., 2009). Between embryonic days 10.5 (E10.5) and 12.5, CR cells migrate tangentially within the preplate to specifically populate the caudo-medial, rostro-medial and lateral cortex (Bielle et al., 2005; Griveau et al., 2010; Villar-Cervino et al., 2013a). CR cell migration and distribution at the surface of the neocortex play important roles in controlling the size and positioning of primary and higher-order cortical areas (Griveau et al., 2010; Barber et al., 2015). It has been proposed that CR cells act as mobile signaling cells that participate in the production and maintenance of morphogen gradients required to coordinate neurogenesis and to delineate areal boundaries within the cortical ventricular zone (VZ)(Griveau et al., 2010).

At the molecular level, transcription factors play key functions in the early regionalization of the cortex (Brunet et al., 2007; O’Leary et al., 2007). Among them, the homeoprotein (HP) family has been the focus of several theoretical and experimental studies showing that they have essential functions in the positioning of boundaries within the developing cortex and, beyond, in the entire nervous system. In particular, Pax6 and Emx2 position the boundary between the primary visual and somato-sensory territories (O’Leary et al., 2007). A link between Pax6 and the control of CR cell migration was suggested by previous studies where accumulation of CR cells in the rostral pallium (Matsuo et al., 1993; Stoykova et al., 2003) and abnormal migration of CH-CR cells was detected in Sey mutant embryos (Ceci et al., 2010). Up to now, the role of HPs in early cortical regionalization was thought to be through intrinsic mechanisms in progenitor cells. However, many HPs, including Pax6, can be secreted and internalized (Prochiantz, 2000; Joliot and Prochiantz, 2004) and thus display a non-cell autonomous function, which has led to the hypothesis that part of their morphogenetic activity might involve their secretion (Holcman et al., 2007; Prochiantz and Di Nardo, 2015; Quininao et al., 2015).

In support of the latter hypothesis, Pax6 was found at the cell surface of ventral neural tube tissues where it regulates the migration of oligodendrocyte precursors (Di Lullo et al., 2011) and the graded expression of extracellular Engrailed, another HP, is a guidance cue for retinal growth cones navigating at the surface of the optic tectum (Wizenmann et al., 2009) (Brunet et al., 2005). We therefore investigated whether Pax6 regulates CR cell migration via a non-cell autonomous activity. To specifically trigger removal of extracellular Pax6 we used a mouse model harboring the sequence for inducible single-chain antibody (scFv) directed against Pax6 in the ROSA26 locus (Bernard et al., 2016). We demonstrate that neutralizing Pax6 in the extracellular space modifies the migration of CH- and S-CR cells.

## Results

### Following CR trajectories in 4 dimensions

By imaging E11.5 flattened cortices in culture, we specifically followed CR cells generated at the level of the CH and septum (CH-CR cells and S-CR cells, respectively) using the Δ*Np73*^*Cre*^ mouse line in which Cre and EGFP are specifically expressed in these two cell populations (Tissir et al., 2009) (Fig. 1A). To compare CR migration in a wild-type environment or upon Pax6 extracellular neutralization (thereafter, mutant), scFvPax6 single chain secreted anti-Pax6 antibody was induced by crossing Δ*Np73*^*Cre*^ and *scFvPax6* mouse lines (Tissir et al., 2009; Bernard et al., 2016). The Pax6 scFv was characterized in earlier studies (Lesaffre et al., 2007; Di Lullo et al., 2011; Bernard et al., 2016). It is specific for Pax6 and devoid of a significant cell-autonomous activity, even when expressed without signal peptide, likely because of the instability of disulfide bonds in the reducing cytoplasmic milieu (Lesaffre et al., 2007).

**Figure 1:**
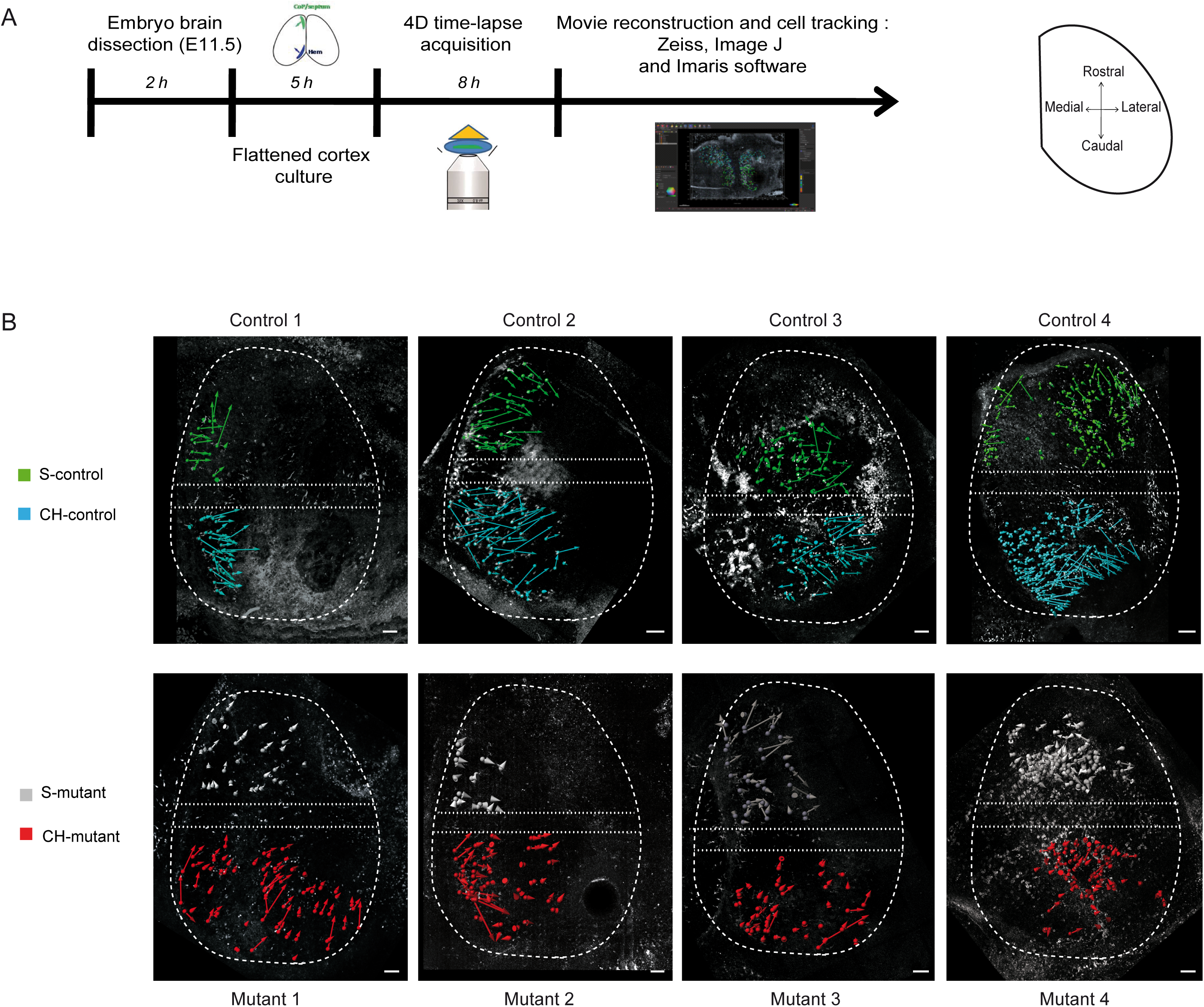
4D-acquisition of CR cell migration patterns. (A) Overview of the successive steps to record and analyze the migration of GFP^+^ CR cells. (B) Cell tracks from 8 hr live imaging of S- and CH-CR cells in E11.5 flattened cortices of Δ*Np73*^*CreIRESGFP*^(control) and Δ*Np73*^*CreIRESGFP*^;*scFvPax6* (mutant) embryos. Tracks are represented by displacement vectors. The CR cells (S or CH) for which origin could not be ascertained were excluded from the images and their localization was delimited by dashed lines. Scale bar: 100 µm.

Four cortices were used for each genotype (wild type and mutant) to track more than 700 CR cells during 8 hrs and obtain 4D views of cell migration (Fig. 1A,B). Initial cell positions were not exactly the same in each cortex, reflecting slightly different embryonic stages (Fig. 1b). Indeed, development proceeds very rapidly and CR cells generated between E10.5 to E11.0 cover the entire pallium within 12 hrs (Bielle et al., 2005; Griveau et al., 2010). Explants with a higher number of medial tracks near the site of origin represent younger stages when migration has just started. Inter-cortical heterogeneity in cell position was observed in control and mutant cortices at the onset and end of filming. We reasoned that if the effect of blocking extracellular Pax6 is robust enough, it should be visible regardless of this heterogeneity and thus we first performed a global analysis before subdividing cortices into several compartments.

### Extracellular Pax6 influences CR cell migration trajectories

To compare CR cell subtypes in the different cortices, we imposed a theoretical migration starting point for all cells in order to analyze each cortex separately (Fig. 2A, B) and to pool the data for each condition (Fig. 2C). We qualitatively confirmed that cell viability was preserved during the 8 hrs of image acquisition by following the progression of migration during 4 successive time intervals in pooled cortices (Fig. S1A). In control conditions for both CR cell subtypes, track path length increased progressively suggesting that cell migration globally continued up to the end of the imaging period. For S-CR cells in mutants, tracks were shorter from the beginning to the end of migration indicating that neurons seemed to cover lower distances than in control conditions. In contrast, tracks of CH-CR cells in mutants evolved similarly to those in controls (Fig. S1A). Furthermore, for both CR cell subtypes, the mean speed after 480 min of recording showed no differences between control and mutant cells suggesting that blocking extracellular Pax6 did not affect their ability to migrate (Fig. S1B). The progressive increase in track length and the maintenance of an average speed over time demonstrate neuron viability in flattened cortices during the entire imaging period.

**Figure 2:**
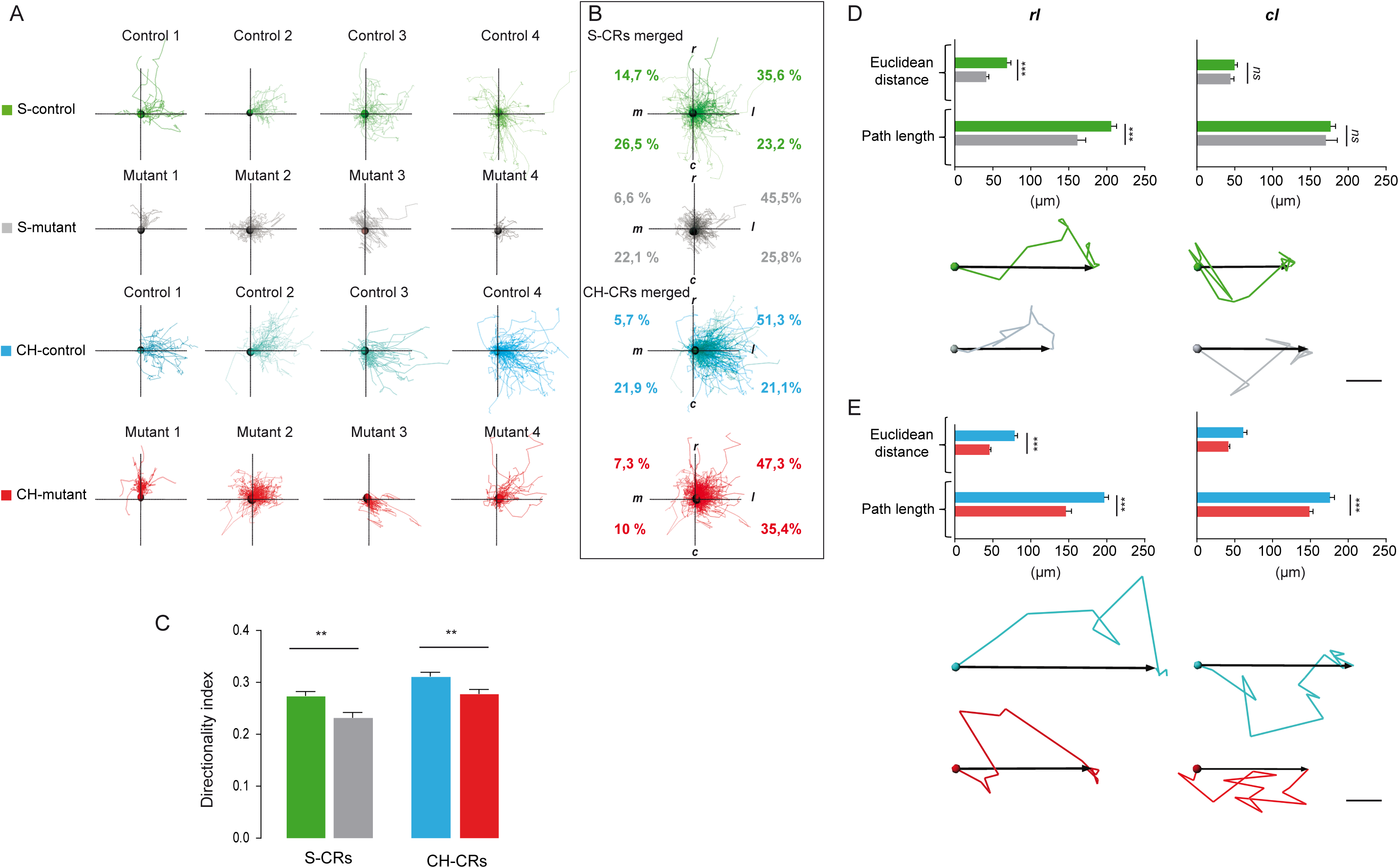
Non-cell autonomous activity of Pax6 on CR cell trajectories and directionality. (A, B) Tracks of migrating S- and CH-CR cells (A) in each control and mutant cortex after 480 min along the medial-lateral and rostro-caudal axes. The tracks are pooled in (B) and the distribution of tracks per quadrant is shown by percentage. (C) Directionality index is the ratio of Euclidean distance to path length trajectory distance. (D, E) Path length and Euclidean distance travelled by CR cells in the RL and CL quadrants. Representative trajectories of S- and CH-CRs are illustrated for each condition. Scale bar: 10 µm. Values are mean ± SEM; ANOVA, *ns*, not statistically different, *\*\*p*< 0.01, *\*\*\*p*< 0.001. CL: caudo-lateral; CM: caudo-medial; RL: rostro-lateral; RM: rostro-medial.

We next analyzed the migration pattern of CR cells after 480 min of recording, in each separate cortex (Fig. 2A) and pooled cortices (Fig. 2B) by quantifying the trajectories along the medio-lateral and the rostro-caudal axis (Fig. 2B). Compared to controls, both mutant S- and CH-CR cells presented shorter track path length with a lower directionality index indicating that mutant CR neurons migrate along less linear trajectories (Fig. 2A-C). The majority of S- and CH-CR cells in controls migrated towards lateral regions with a preference for the rostro-lateral (*rl*) direction (35.6% and 51.3 %, respectively) (Fig. 2B). In mutants, compared to controls, the percentage of S-CR cells migrating in *rl* directions increased (45.5% *vs* 35.6%, respectively) (Fig. 2B) whereas that of CH-CR cells migrating caudo-laterally (*cl*) was preferentially enhanced (35.4% *vs* 21.1%, respectively) (Fig. 2B). To further examine these differences, we analyzed distances travelled by CR cells focusing on lateral directions (Fig. 2D, E). Extracellular Pax6 neutralization was efficient in decreasing path length of S-CR cells migrating rostro-laterally (39.73%) with a smaller but significant effect on Euclidean distance (21.61%). Notably, no differences were observed for S-CR cells migrating caudo-laterally. For CH-CR cells, Pax6 neutralization resulted in a reduction of both path length and Euclidean distance values whether they migrated *rl* or *cl* (25.59% *vs* 41.82% and 15.34% *vs* 32.02%, respectively). A representative trajectory of CR cells in each condition (Fig. 2D, E) illustrates that neutralizing extracellular Pax6 modifies the migration of S-CR and CH-CR neurons toward the lateral part of the developing cortex with a stronger decrease in overall Euclidean distance than total path length. Together these results suggest that secreted Pax6 influences more strongly directionality than the ability to migrate.

### Position-dependency of non-cell autonomous Pax6 activity

To study whether the effect of extracellular Pax6 neutralization depends on the region of the developing cortex where CR cells migrate, we took into account the differences in CR cell track start positions. Cortices were subdivided into 4 compartments along the medio-lateral and the rostro-caudal axis with cell numbers ranging from 97 to 265 per compartment to allow for statistical analyses. The two rostral territories (RM and RL) are enriched in S-CR cells, while the caudal ones (CM and CL) primarily contain CH-CR cells. In addition, RM and CM compartments are closer to the CR cell sources than RL and CL compartments which abut the “arrival regions”. CR migration was analyzed separately for the 4 compartments (Fig. 3A-D). In all 4 compartments both S- and CH-CR cells in mutants traveled shorter Euclidean distances along the medio-lateral axis with respect to controls. However, along the rostro-caudal axis migration of both CR subtypes was only influenced by Pax6 neutralization in the lateral compartments where both S- and CH-CR cells appeared to travel shorter Euclidean distances and to move less along the rostro-caudal axis (Fig. 3C, D). The mean directionality vectors of controls and mutants (Fig. 3E) summarize the effect of blocking extracellular Pax6 on trajectories in the different territories. They show that distances traveled are reduced for mutant CR cells in all 4 compartments while the direction of migration is more highly affected in the lateral cortex with an overall increase of migration in the rostral direction at the expense of caudal direction. Taken together, these results show that neutralization of extracellular Pax6 reduces the distance traveled by CR cells leaving their medial origins as well as when reaching the end of migration in the lateral cortex. Notably, they also indicate that in the lateral compartments, where the Pax6 concentration is higher rostrally, the neutralization of its extracellular activity redirects the cells in the rostral direction, suggesting a repelling activity of high extracellular Pax6 concentration.

**Figure 3:**
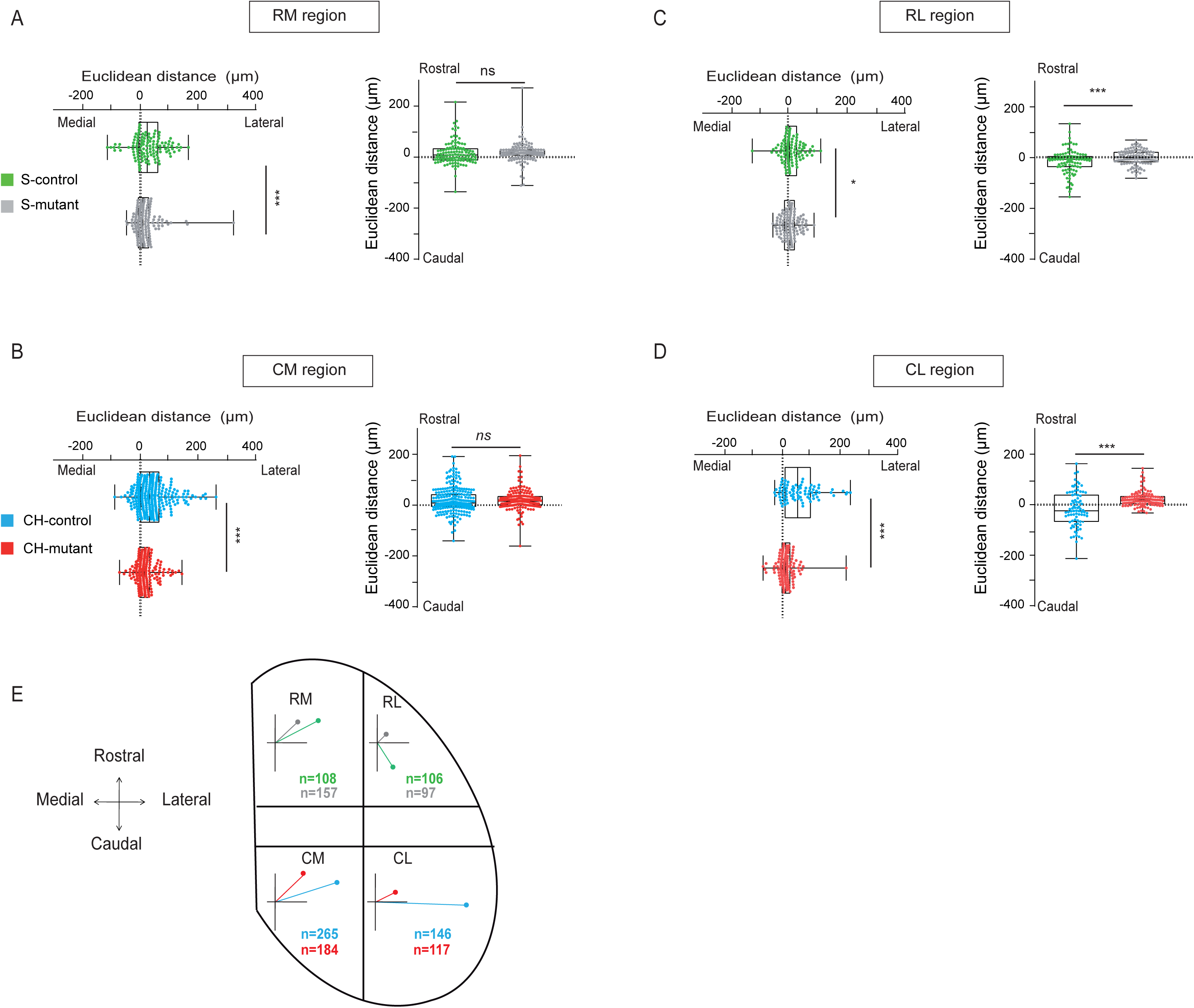
Position-dependency of non-cell autonomous Pax6 activity. (A-D) S- and CH-CRs migration (Euclidean distance) in the 4 compartments: RM (A), CM (B), RL (C) and CL (D) for control and mutant cortices. Each point represents a cell. (E) Control and mutant cortices were subdivided in 2 sections along the medio-lateral axis and in 3 sections along the rostro-caudal axis. In the 4 select compartments, mean displacement vectors were calculated from CR cell tracks. Vectors represent the Euclidean distance travelled during 8 hrs of migration. Values shown are mean ± SEM; ANOVA, *ns* not statistically different, *\*p*<0.05, *\*\*\*p*< 0.001.

### Blocking extracellular Pax6 affects the depth of CR cell migration

Previous studies have analyzed migration parameters of CR cells in 2-D or 3-D environments (Villar-Cervino et al., 2013a; Barber et al., 2015). Migration within the depth of the tissue has never been analyzed in whole-flattened cortical preparations because of the excessive size of the movies. We have now devised a method to analyze over 15 GB time-lapse files by movie frame separation and reconstruction (see Methods). We were thus able to quantify the effect of Pax6 neutralization on S- and CH-CR localization within the depth (Z axis) of the developing cortex as a function of time in the 4 compartments of interest (Fig. 4). Although percentages along the Z-axis might not represent real in vivo values due to tissue shrinking in flattened cortex cultures, our data demonstrate that close to CR sources (RM and CM domains), S- and CH-CR cells migrated at 40% or 30% depth respectively, with a wider dispersion between the ventricular side and the pial surface in the mutant CH-CR cells (Fig. 4A). When closer to the end of their migration (RL and CL territories), CH-CR cells in controls appeared to spread highly along the depth compared to S-CR cells which migrated at about 50% depth. This behavior was lost in the mutant environment where both S- and CH-CR cells migrated close to the pial surface (Fig. 4B). Representative images of the different regions of interest (Fig. 4A,B) demonstrate that blocking extracellular Pax6 strongly changes the distribution of CR neurons along the depth of the developing cortex.

**Figure 4:**
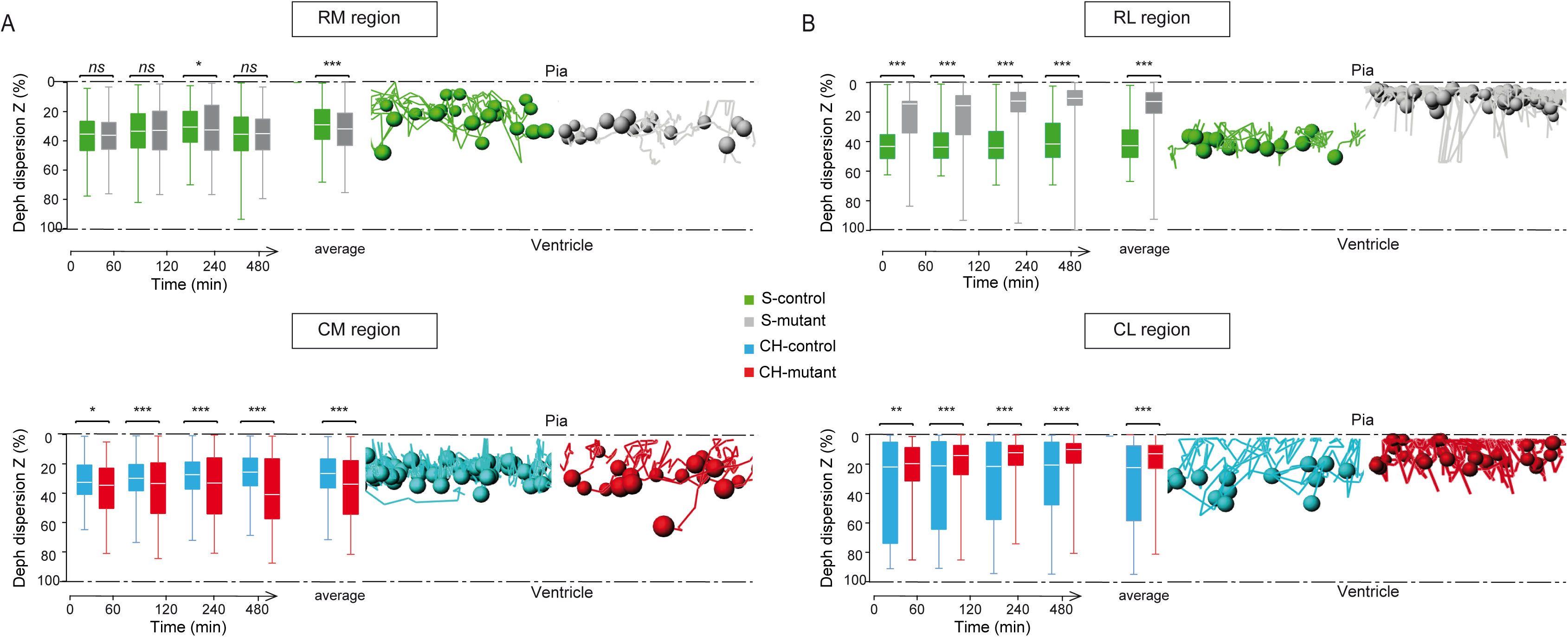
Blocking extracellular Pax6 affects CR cell depth. (A, B) Position of S-(A) and CH-CR (B) neurons in the depth axis (Z) for control and mutant cortices, classified in compartment at the onset of tracking (RM, RL, CM and CL). For each cortex, values were weighted for comparison and are represented by percentage. CRs localization is illustrated by representative S- and CH-CR cells in each condition. Values shown are mean ± SEM; ANOVA, *ns*, not statistically different, *\*p*<0.05, *\*\*p*< 0.01, *\*\*\*p*< 0.001.

### Non-cell autonomous effects on CR cell contact repulsion

CR cells, in particular CH-CR cells, were shown to tangentially invade the developing cortex by contact-repulsion: when the growth cone of a CR cell comes into contact with another CR cell, its leading process retracts, its direction of migration changes by more than 90°, and its migration speed increases (Villar-Cervino et al., 2013a). In support of contact-repulsion, we found that CR cells migrate faster at the onset of migration when cell density is higher and, in particular, CH-CR cells displayed enhanced migration speed upon neutralization of extracellular Pax6 (Fig. S2A, B). After 1hr of recording, the speed was reduced for both CR cell subtypes with lower values recorded in mutants (Fig. S2A, B) suggesting that blocking extracellular Pax6 may have contrasted effects at the onset and at the end of migration.

We therefore tested whether changes in contact-repulsion may be involved in the migration defects observed in mutants (Fig. 5). GFP^+^ CR cells were clustered into two groups depending on whether or not they encounter and contact another GFP^+^ CR cell during migration (Fig. 5A). In control conditions, S- and CH-CR cells behaved differently, as CH-CR cells displayed higher number of contacts in medial and lateral domains compared to S-CR cells (74% in CL and 71% in CM *vs* 64% in RM and 56% in RL). Blocking extracellular Pax6 increased contact numbers between S-CR cells in the RL domain (73% *vs* 64%) and, on the contrary, decreased contact numbers in CH-CR cells in both CM and CL territories (Fig. 5A). In a more thorough analysis, we further separated cells initiating contacts in two groups (Fig. 5B-D): *(i)* contact-repulsion (Fig. 5B); and *(ii)* contact-without-repulsion when following an encounter, cells remain in contact longer and do not change their direction of migration (Fig. 5C). In control conditions, a higher percentage of both CR cell subtypes showed contact-repulsion in medial than in lateral domains confirming previous results that contact-repulsion could be an important mechanism for cell dispersion near CR sources (Villar-Cervino et al., 2013b). Blocking extracellular Pax6 enhanced contact-repulsion in medial regions and significantly diminished it in lateral regions (Fig. 5D). These differences were more marked for CH-CR cells suggesting that neutralization of Pax6 is more efficient in caudal regions.

**Figure 5:**
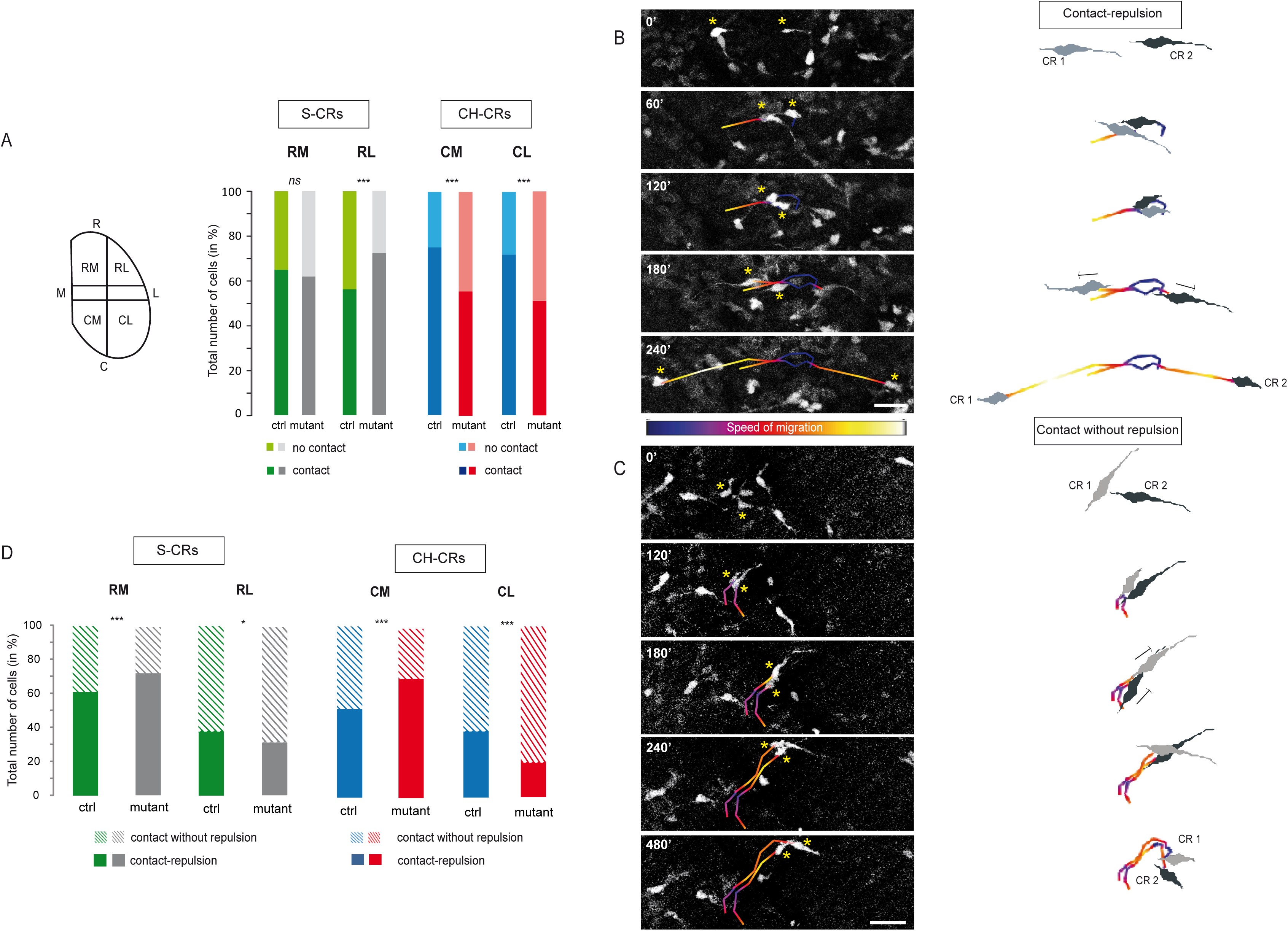
Pax6 neutralization stimulates or inhibits CR cells undergoing contact-repulsion in a regions-dependent manner. (A) S- and CH-CR cells were clustered by: *contact* if a CR cell encounters another CR cell during migration; *no contact* if a CR cell migrates without contacting another GFP^+^ positive cell. (B-D) Contact between two cells was subdivided in two categories: (B) *contact-repulsion* when contact with another CR cell resulted in the full retraction of their leading process, re-extension of a new leading process and change of initial direction (images taken from control CH-CR cells); (C) *contact-without-repulsion* when contact with another CR cell did not change the direction (images taken from mutant S-CR cells). Tracks are color coded for speed of migration. Directionality of migration is represented by a white arrow. (D) Quantification of contact-repulsion (filled bars) and contact-without-repulsion (hatched bars) of S- and CH-CR cells in control and mutant cortices. Values shown are mean ± SEM; ANOVAt, *ns*, not statistically different, *\*p*<0.05, *\*\*\*p*< 0.001. Scale bar: 50 µm.

## Discussion

In this study we used a genetic strategy to trigger the expression and secretion of a single chain antibody against Pax6 to neutralize it in the extracellular space in the developing cortex in order to study its non-cell autonomous function during neuronal migration. We show that neutralizing this secreted transcription factor modifies the tangential and Z trajectories and migration behavior of S- and CH-CR cells. This genetic strategy validated earlier for Pax6, Engrailed and Otx2 (Lesaffre et al., 2007; Wizenmann et al., 2009; Di Lullo et al., 2011; Layalle et al., 2011; Bernard et al., 2016) decreases both extracellular Pax6 and Pax6 internalization. Based on these previous results, we favor the idea that non-cell autonomous Pax6 activity requires its internalization to cause changes at the transcriptional, translational and epigenetic levels. However, we cannot preclude a pure extracellular effect triggered by Pax6 extracellular binding sites or receptors. A profound understanding of the mechanisms involved requires further study at a molecular and cellular level.

Characterizing the factors which control correct positioning of CR neurons at the cortical surface is a crucial issue since perturbing CR subtype distribution and migration alters the position and size of primary (Griveau et al., 2010) and higher-order areas (Barber et al., 2015). Using live imaging and genetic labeling, we followed S- and CH-CR cells during early phases of migration. Upon extracellular Pax6 neutralization, we observed similarities and differences in the responses of the two CR subtypes. In terms of path length and Euclidian distances, neutralizing Pax6 had a negative effect, although stronger on Euclidian distance, both on S-CR and CH-CR cells migrating with a *rl* direction towards the high concentration of Pax6 gradient expression. Notably, this had no effect on S-CR cells whose tracks were oriented along the *cl* direction away from the high Pax6 concentration. In terms of directionality, S-CR and CH-CR cells responded similarly with a strong tendency to migrate more rostro-laterally in the mutant, an effect particularly striking for S-CR cells in the lateral region where Pax6 expression is higher. Together these results suggest a repulsive activity of high extracellular Pax6 concentration in the most lateral regions. Furthermore, given that for both subtypes the effect of Pax6 neutralization on Euclidian distance is stronger than on path length and that a reduction in directionality index is observed without changing average migration speed (Fig. S1), the phenotype reflects a disorganization of migration with little effect on the ability to migrate. Wild-type CH-CR cells were previously found to migrate by expanded routes in *Sey* mutants embryos, in which both intracellular and extracellular Pax6 are eliminated, compared to control embryos (Ceci et al., 2010). They appeared to travel longer distances and to disperse more broadly along the rostro-caudal and Z axis. Interestingly, we observed that CR cells of both subtypes travelled shorter distances and dispersed less, on both axis, upon specific removal of extracellular Pax6. This difference is best explained by the fact that, in contrast with the *Sey* mutant situation, neutralizing Pax6 non-cell autonomous signaling functions leaves intact its cell autonomous transcriptional activities, suggesting a specific function of secreted Pax6 on CR cell migration. We also find that the effect on CR cell directionality upon neutralization of secreted Pax6 depends on cell position at the onset of recording and that migration direction along the rostro-caudal axis is more highly disturbed in lateral regions, where Pax6 is expressed at higher levels. This dose- and region-dependent activity suggests that the graded intracellular Pax6 distribution in the embryonic cortex is preserved in the extracellular space and that secreted Pax6 is poorly diffusible. Indeed, extracellular HPs, as also found for several classical morphogens (Prochiantz and Di Nardo, 2015), are unable to diffuse over long distances when not within complexes, and thus remain bound to the cell surface, probably trapped in the local extracellular matrix. This was confirmed in the case of Engrailed in the optic tectum (Wizenmann et al., 2009) or in the Drosophila wing disk (Layalle et al., 2011), and in the case of Pax6 in the ventral neural tube (Di Lullo et al., 2011).

The analysis of CR cell dispersion within the neuroepithelium (Z axis) reveals some differences in the effect of Pax6 neutralization between S-CR and CH-CR cells and also between territories. In the medial compartments (RM and CM), neutralizing Pax6 had little influence on migration in the Z axis, although a wider dispersion was noticed for CH-CR cells in the mutant. In the lateral compartments, both S-CR and CH-CR cells migrated closer to the surface in the mutant than in the wild type. This effect was milder for CH-CR cells since control cells navigated more randomly dispersed along the Z axis. It has been proposed that chemokines such as CXCL12 secreted by the meninges both enhance CR cell motility and maintains them close to the pial surface (Borrell and Marin, 2006). Our results thus suggest that extracellular Pax6 and CXCL12 may have opposite effects on the migration of CR cells in the Z axis and on their maintenance in the marginal zone.

Contact-repulsion was shown to be an important mechanism by which CR cells populate the cortex at early stages. In live-imaging studies, CR cells are repelled upon contact with neighbor cells resulting in the collapse and retraction of their leading process and a change in their direction of migration (Villar-Cervino et al., 2013b; Barber et al., 2015). It was also shown that a proportion of neurons randomly change their migration direction without contacting adjacent cells (Villar-Cervino et al., 2013b). In the case of S-CR cells, neutralizing Pax6 increased the number of contacts in the RL but not in the RM compartment, although contacts were more often followed by repulsion in the latter compartment. In the case of CH-CR cells, we found a decrease in the number of contacts in the CM and CL with increased and decreased repulsion in the CM and CL compartments, respectively. These results suggest that there is no systematic correlation between contact and repulsion. Eph/ephrin signaling was previously shown to be involved in contact-repulsion. Abrogating this pathway increased the duration of CH-CR cell contact and their persistence in directionality, thus reducing their ability to undergo contact-repulsion (Villar-Cervino et al., 2013b). Pursuing this hypothesis will require full analysis of the molecular mechanisms involved in Pax6 activity. Nevertheless, we underscore the similarity between CR cell guidance by Pax6 and retinal growth cone guidance by Engrailed, which raises the possibility that, similarly to what was demonstrated for Engrailed (Wizenmann et al., 2009), Pax6 co-signals with Ephrins and/or other bona fide morphogens, including chemokines.

Altogether, our findings show for the first time that extracellular Pax6 controls non-cell autonomously CR neuron migration, an observation analogous to the paracrine activity of Pax6 in oligodendrocyte precursor cell migration in the embryonic chick spinal cord (Di Lullo et al., 2011). This supports the hypothesis that several mechanisms can be employed by a gene/protein to execute the same function. We suggest that Pax6 may control cortical arealization by directly acting as a transcription factor in neuronal progenitors and indirectly as a guidance factor for proper CR cell migration. Therefore, slight changes in the levels and distribution of extracellular Pax6 during brain development could have important consequences in physiopathology and be at the origin of evolutionary modifications in the mammalian cerebral cortex.

## METHODS

### Animals

*scFvPax6* (Bernard et al., 2016) and Δ*Np73*^*CreIRESGFP*^(Tissir et al., 2009) transgenic mice were kept in a C57BL/6J background. Animals were genotyped by PCR using specific primers for each allele. The day of vaginal plug was considered to be E0.5. All animal procedures, including housing, were carried out in accordance with the recommendations of the European Economic Community (86/609/EEC), the French National Committee (87/848) and French bylaws (AGRG1240332A / AGRG1238724A / AGRG1238767A / AGRG1238729A / AGRGR1238753A). All mouse work was approved by the Veterinary Services of Paris (Authorization number: 75-1454) and by the Animal Experimentation Ethical Committee Buffon (CEEA-40) (Reference: CEB-34-2012).

### Whole flattened cortices culture

Whole flattened cerebral cortices cultures were adapted from Barber M., *et al* 2015. Cortices were dissected from E11.5 Δ*Np73*^*Cre/+*^(control) and Δ*Np73*^*Cre/+*^;*scFvPax6*^*tg/o*^(mutant) embryos. One single incision was made to remove the ventral subpallial tissue. Explants were cultured for 5 hours (hrs) before time lapse acquisition on a Millicell permeable membrane (pore size 0.4 µm; Millipore) with the ventricular side facing the membrane, in phenol-red-free high glucose DMEM (Sigma) containing B27 supplement (Gibco) and penicillin/streptomycin antibiotics (100 µg/ml, Sigma) in 5% CO_2_ at 37°C. Shortly before imaging, the flattened cortex was transferred with the Millicell membrane and inverted onto a glass-bottom microscope imaging chamber with the pial surface directly adjacent to the glass. Explants were mounted in purified bovine collagen gel (25 µl/explant, Advanced Biomatrix), which polymerizes in 15 min at 37°C.

### 4D time-lapse acquisition

Time lapse acquisition of migrating CR cells was performed over 8 hrs in an incubation chamber at 37°C and 5% CO_2_ in phenol red-free DMEM solution by using an inverted laser scanning confocal microscope (Zeiss 780) with an oil immersion 25x objective. EGFP protein was excited at 488 nm and images were acquired at 500-550 nm. Depending on explant size, 4D time-lapse images were generated by acquiring automatically 9-14 optical Z sections with a z-step size of 8 µm every 30 min and 70-140 focal tiles were merged to visualize the entire flattened cortex with a 512 × 512 pixel resolution.

### Bioinformatics processing and movie analysis

Successive steps were used to disassemble and divide resulting movies, exceeding 15 GB, in time and Z points successively by using the Zeiss Zen software (Carl Zeiss MicroImaging) and the ImageJ software to convert images in 3D tiff-stacks. 4D reconstruction and display of CR cell migration were built up by Imaris software (Bitplane, version 8.2.1). GFP^+^ cell tracks were obtained from four flattened brains per genotype (control or mutant). Using the spot cell-tracking module, cells were spatially classified depending on their genotype and origins (CH or S) and tracked manually over time and along the medio-lateral, rostro-caudal and depth axes. We selected CR cells migrating in the caudal quadrant of the explant as corresponding to CH-CRs cells, and those migrating in the rostral quadrant as S-CR cells (according to (Barber et al., 2015)). We excluded from the analysis CR cells migrating at middle levels along the rostro-caudal axis for which the origin could not be ascertained. 1478 tracks were analyzed, 726 control cells and 752 mutant cells. The directionality index was calculated by dividing the Euclidean distance (the shortest distance between the start and end position of a cell trajectory) by the actual path length followed by the cells during migration. The overall directionality in migration was determined for each region of interest by automatic clustering tracks to a common origin using the Imaris transpose tracks function. In Figure 3E, a mean displacement vector along the medio-lateral and rostro-caudal axis was calculated for each region and represented in a 2-D vectorial projection clustered at a common origin. CR cell behaviors at contacts were subdivided and analyzed based on previous strategies (Villar-Cervino et al., 2013b; Barber et al., 2015). Briefly, contacts between two cells were scored if occurring within a single optical slice from Δ*Np73*^*Cre/+*^ (control) and Δ*Np73*^*Cre/+*^;*scFvPax6*^*tg/o*^ (mutant) explants in the rostral and caudal regions. The angle of deviation was observed using the Imaris cell-tracking spot module and considered contact with repulsion if cells changed trajectory with an angle of more than 90°.

### Statistics

Prism 6 software (GraphPad, version 6.01) was used for statistical analysis. When data followed a normal distribution, paired comparisons were analyzed with t test, whereas multiple comparisons were analyzed using one-way ANOVA with post hoc Bonferroni correction. Data are presented as mean and ±SEM throughout the manuscript. p < 0.05 were considered significant.

## Author contributions

H.K. designed and performed most of the experiments, A.W. participated in several experiments, M.V. and A.A.D. produced and validated the *scFvPax6* mouse line, E.C. helped analyzing the data. A.P. and A.P. supervised the project and analyzed data. H.K., E.C., A.A.D., A.P. and A.P. wrote the manuscript.

## Acknowledgements

We thank the ImagoSeine facility, member of the France BioImaging infrastructure supported by the French National Research Agency (ANR-10-INSB-04, “Investments for the future”), for help with confocal microscopy and Imaris software. We thank the Buffon Animal facility, Animalliance and L. Vigier for technical assistance and animal care. F. Tissir for providing the Δ*Np73*^*CreIRESGFP*^ mouse line (Brussels, Belgium). F. Lam and V. Contremoulins for help with movie reconstruction. A. Pierani is a CNRS (Centre National de la Recherche Scientifique) Investigators and member Teams of the École des Neurosciences de Paris Ile-de-France (ENP), E.C. is a University Paris Diderot Lecturer. This work was supported by grants from the ANR-MOST bilateral accord (ANR-13-ISV2-0001) to A. Prochiantz and A. Pierani, ERC Advanced Grant HOMEOSIGN (339379) to A. Prochiantz, ANR (ANR-2011-BSV4-023-01; ANR-15-CE16-0003-01) and FRM («Equipe FRM DEQ20130326521») to A. Pierani.

## Data availability

**Figure S1:**
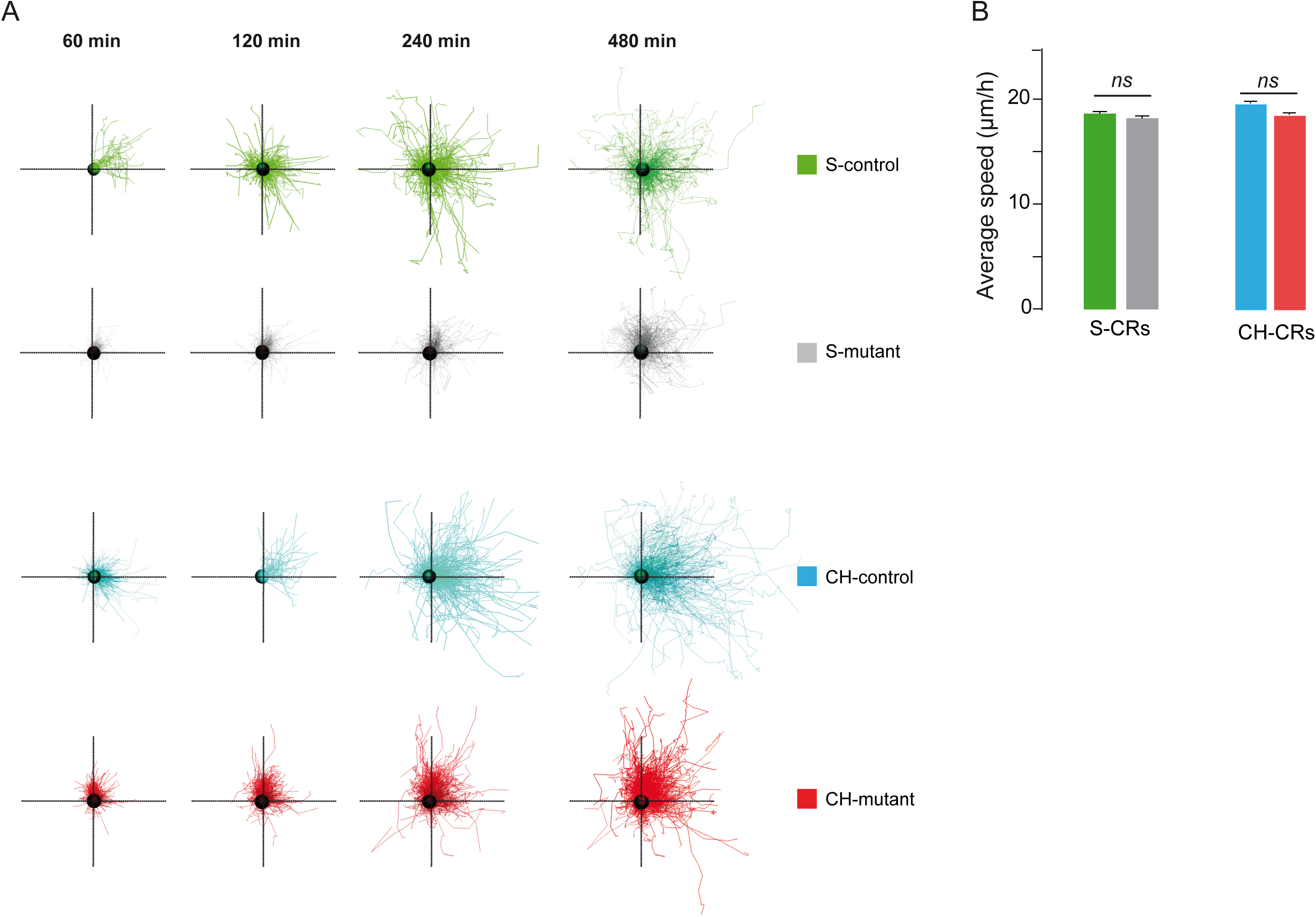
Migration trajectories and speed at different time points. (A) Trajectories of S- and CH-CR cells at different time points. A theoretical starting point was taken and all cells from the 8 cortices (4 control and 4 mutant cortices) were pooled. (B) Average speed of S- and CH-CR cells in controls and mutants after 480 min of recording. All cells from the 8 cortices were analyzed. Values shown are mean ± SEM; ANOVA, *ns* not statistically different.

**Figure S2:**
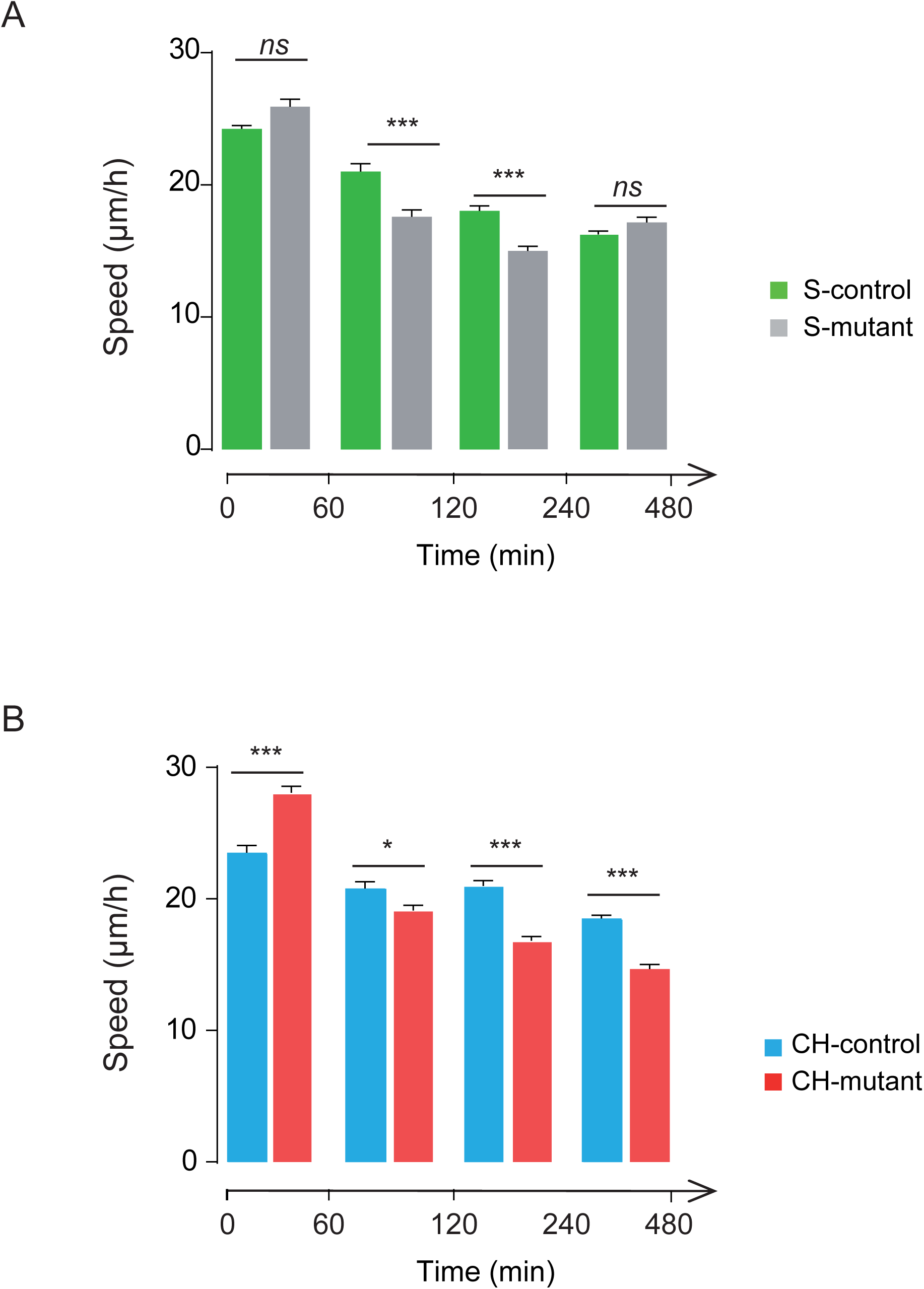
Analysis of migration speed of S- and CH-CRs (related to figure 5) Speed of S-(A) and CH-CR (B) cells in controls and mutants were evaluated at different time points. All cells from the 8 cortices were analyzed. Values shown are mean ± SEM; ANOVA, *ns* not statistically different, *\*p*<0.05, *\*\*p*< 0.01, *\*\*\*p*< 0.001.

